# Cool Enough for School: Second Version of Google Glass Rated by Children Facing Challenges to Social Integration as Desirable to Wear at School

**DOI:** 10.1101/171033

**Authors:** Ned T. Sahin, Neha U. Keshav, Joseph P. Salisbury, Arshya Vahabzadeh

## Abstract

**Background:** On July 18^th^, 2017, X, a subsidiary of Alphabet Inc. announced the successor to Google Glass. Glass Enterprise Edition could function as an assistive technology for autism spectrum disorder (ASD), yet its acceptability, desirability, and the willingness of children with ASD to wear it, are not known. The authors review key issues surrounding smartglasses and social communication, child development, and the school environment.

**Methods:** The smartglasses were evaluated by eight children with ASD, who ranged from moderately non-verbal to verbal, in the context of whether they would desire to wear it and use it as an assistive device each day at their respective schools. Children represented the full range of school ages (6 – 17yrs).

**Results:** All eight children responded that they would want to wear and use Glass Enterprise Edition at school. Additionally, all eight parents said their child had fun during the testing session, and six of eight parents said the experience went better than they had thought.

**Conclusion:** The results show that children with ASD are willing to use Glass Enterprise Edition in a school setting, highlighting its desirability and social acceptability in this population, as well as its future potential as an assistive technology.

## INTRODUCTION

Autism Spectrum Disorder (ASD) is a childhood onset developmental disorder with a rapidly rising prevalence, with 3.5 million people with ASD in the United States alone (1). Innovative assistive technologies may help to address the unmet educational and therapeutic resource demands of the ASD community (2). While there are many different types of assistive technology, the portability, capability, and ubiquity of smartphone and tablet devices has led to considerable growth in assistive apps for these devices (3, 4). More recent technological advances have led to the development and release of a range of smartglasses, face-worn computers with a visual display, and a range of in-built sensors (5–7).

Smartglasses are capable of delivering a variety range of experiences, including augmented and virtual reality (8). They are sensor-rich, and are able to collect an extensive range of quantitative user data (9–12). This data can be monitored and analyzed on a real-time basis, allowing for the smartglasses to dynamically change the user experience to optimize learning, effectively placing the user and the smartglasses in a closed feedback loop (13, 14). Given the proximity of smartglasses to the sensory components contained in the human head, this type of computing will enable a higher level of human-computer interaction (14). Smartglasses are already being developed as a social and behavioral communication aid for people with ASD (13, 15–17). There are also a number of differentiating factors worthy of consideration when we compare handheld devices to smartglasses. Hand-held devices such as tablets and smartphones require one or both hands to hold the device, and encourage a heads-down posture (Figure 1a, left) (18). Evidence suggests that smartphone use may decrease users’ awareness of their social and physical environment; this is a particular concern in people with ASD, given that they already face challenges engaging with the social world around them (19). In contrast, head-worn computers pose an advantage in allowing and potentially encouraging children to remain heads-up while using them. This gives users the ability to better engage with the social world while using head-worn computers, while interacting with classmates or parents, for example (Figure 1a, right).

**Figure 1.**
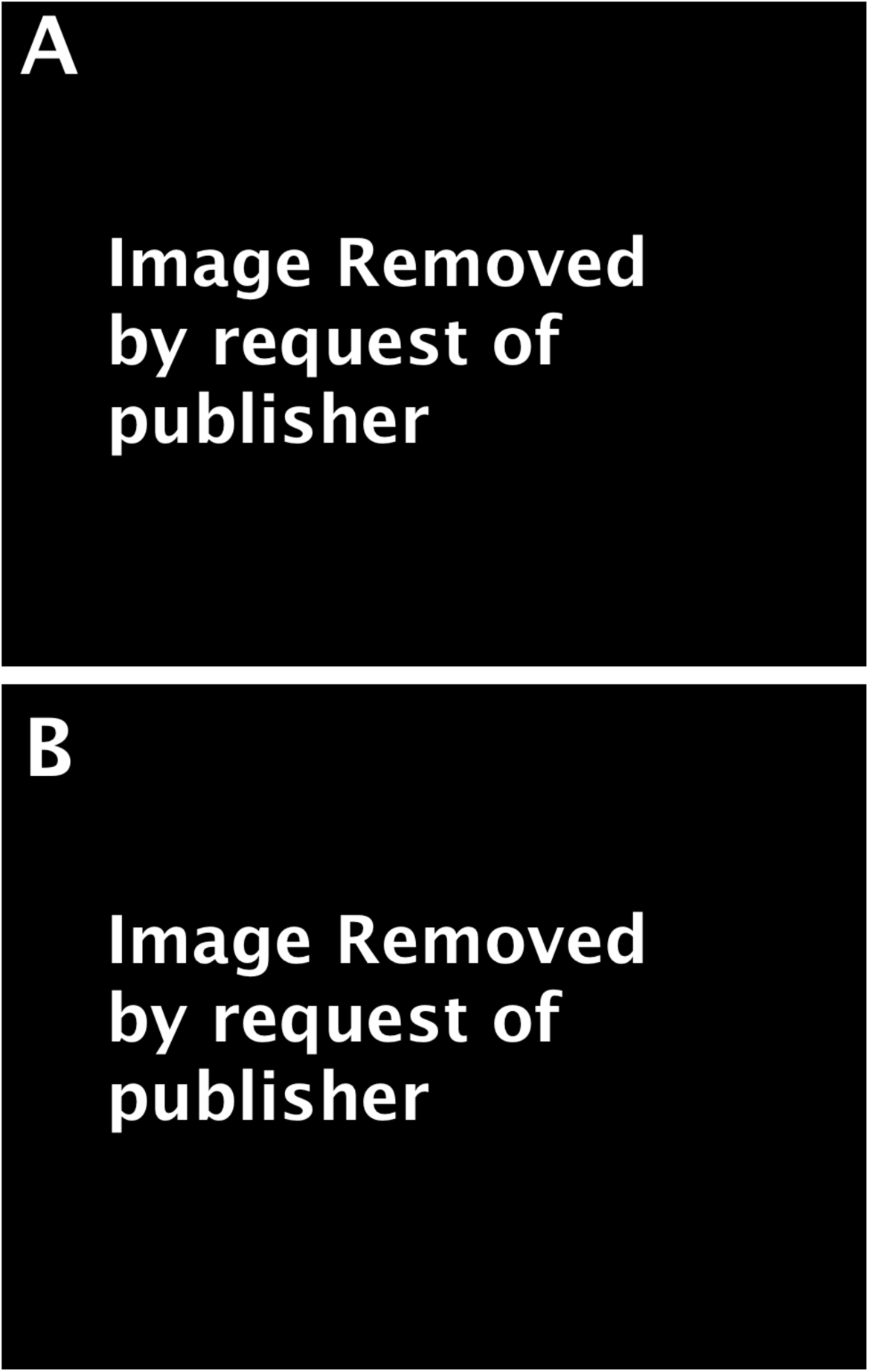
Head-worn computers encourage users to be heads-up and allow them to be hands-free, in contrast to screen-based technologies such as phones and tablets. A.) Demonstrative example of a person using a tablet while her sibling uses Glass Enterprise Edition. Tablet use encourages a heads-down stance, suboptimal posture, and visual disconnection from the social world. B.) The Glass EE device from multiple views.

There have only been a handful of reports on the use of smartglasses in people with ASD (13, 15–17), with little research on their attitudes towards using and wearing these relatively novel devices. The use of smartglasses in people with ASD also requires discussion of their potential impact on social communication from a cognitive neuroscience standpoint, and their prospective influence on child development from ecological, psychosocial, and cognitive child development theories. The personal desires of ASD children and young adults are crucially important, because they are the intended users and beneficiaries, and especially because children with challenges or different types of minds are often forced to use devices and systems they do not actually like, or want to be associated with (20, 21). This is ultimately less effective because aversion leads to lower compliance. Poor adherence and problems with maintaining lasting engagement are some of the largest issues facing educational devices and applications, as well as well-being and lifestyle tools (22, 23).

Many people with ASD use assistive technology to help them with communication skills, social and emotional skills, and adaptive/daily living skills (24). Assistive technologies elicit a range of responses, and can be considered “cool” (21), “weird”, desirable, or a source of stigma (25, 26). Users of assistive technologies can often express a preference for the type of assistive technology that they want to use (27, 28), even at a young age (29). Additionally, the social acceptability of an assistive technology may be one of the most important elements in determining if that technology is used by people with developmental disabilities (26, 30). These individuals have often had to use technologies that have been selected for them and their families while having little input to the potential negative image, stigma, or embarrassment in using such technologies (26). Understanding and implementing user preference of assistive technologies empowers self-determination in these individuals (27). The preferences and views of the family and caregivers of these individuals are also important as they impact the acceptance and effective use of such technologies in the household (24, 31). These issues are clearly pertinent to smartglasses, especially in light of the multiple reports of negative public perception of these technologies, mostly around privacy concerns (32).

### Potential impact of smartglasses on social communication

The human face, a complex and dynamic system, is our most powerful means of social communication (33). To successfully transmit social information to another person, the sender must have the mental and physical means of generating a facial and bodily representation of the social information that they wish to send, while the receiver must be in a position to see and decode the facial and bodily representations into social information. The social communication deficits seen in ASD may impede the ability to both send and receive social information. People with ASD are reported to have deficits in facial perception (34, 35), emotion recognition (36), eye gaze (37) and production of facial expressions (38). It is important to consider the possibility that social communication may be further impacted by the physical presence of smartglasses on a sender’s face. Smartglasses may impede social communication if, for example, the sender demonstrates a hesitancy in producing natural head movements or expressing large magnitude facial emotional expressions due to concern that the smartglasses may fall off the face, or be damaged. Smartglasses may also impair social interaction if the user feels the assistive device is socially undesirable (39) or a source of stigma (26). In these situations, the user may not use the device, or may alter their facial and bodily actions to minimize attention to themselves. Furthermore, the physical form factor of smartglasses may obscure a portion of the wearer’s face that is visible to others, especially the central information-rich parts of the face, such as the eye regions (40). The effect of this may be dependent on the age of the sender, as biologic age determines human head size (41) and therefore the portion of the face that would be obscured. It may also depend on the ability of the receiver to successfully compensate for partly missing facial data and to make inferences about a sender (a common application of this in ASD research is the “Reading the Eyes in the Mind” test (42)). Since both people with and without ASD find it difficult to read the facial emotional expressions of people with ASD (38), it is conceivable that further obscuring the amount of visible facial information could make the interaction more arduous. This point may be particularly relevant to interactions between people with ASD and their unaffected family members. ASD is a highly hereditable condition with a complex genetic basis (43), and many unaffected relatives of children with ASD have been found to have subclinical autistic traits (44).The parents of children with ASD may demonstrate subtle deficits in social communication and face processing (45, 46). It may be sensible to minimize facial obscuration given that the aim of assistive smartglasses is to enhance social communication between people with ASD and their family members.

The presence of face-worn smartglasses may also influence social relationships as they alter a user’s facial appearance, and unlike many other assistive technologies, they are not easy to hide. Wearing smartglasses may not only alter how the user perceives the world, but may alter how the world perceives the user. Facial appearance plays a key role in determining how people interact with one another (47), including who they help, hire, or want to date (48). Human faces may also be judged based on their symmetry, a marker of attractiveness and an indicator of optimal developmental outcome despite environmental stressors (49). Greater facial symmetry has been linked to increased perceived trustworthiness, and a decreased risk of bullying (50). Facial symmetry may be perceived as demonstrating genetic quality, and therefore suitability of an individual as a mate (49), while facial asymmetry may be an indicator of psychological, emotional, and physiological distress (51). Users of smartglasses that are asymmetrical, such as those that are monocular, could be perceived as being less attractive and trustworthy due to these evaluative evolutionary mechanisms. By extension, “asymmetric” smartglasses users may also be at greater risk of bullying (50). Yet on the other hand, smartglasses that are asymmetrical may obscure less of the wearer’s face from the view of others. As discussed earlier, maximizing how much of the face is visible may help facilitate social communication. Even non-technological face-worn glasses are associated with impaired interpersonal relationships: for example, wearing prescription glasses or having a history of using eye patches has been associated with a 35% increase in the likelihood of physical or verbal bullying (52).

### Smartglasses in the context of child development

The perceptual impact of smartglasses and their ability to augment a child’s cognitive and emotional functioning may have a central and influential role in childhood development if we consider Bronfenbrenner and Ceci’s *bioecological model* (53) and Brofenbrenner’s earlier *ecological systems theory* (54). According to *bioecological model*, children are active participants in their environments, and they have unique bidirectional interactions with each of their contextually separate environments, including home and school. This model places increased emphasis on the cognitive, emotional, and physical attributes of the child in their development, and in how the child and environments interact with one another. As outlined in Brofenbrenner’s *ecological systems theory* (54), the school environment, like the home environment, is one of the most intimate and influential environments affecting their childhood development, as it lies in the child’s *microsystem*. When we consider that smartglasses may enhance the cognitive and emotional functioning of children within their *microsystem*, we can see that they may have a highly influential role in child development. Even within the *microsystem*, the contextual differences between the most intimate of environments may affect a child’s view towards using assistive technology. Research has shown that children have different attitudes and levels of enthusiasm towards using assistive technology depending on whether they are asked to use it at home or at school (28).

Furthermore, use of smartglasses by future school-age children and adolescents should prompt a discussion of Erikson’s *4^th^ and 5^th^ psychosocial stages* (55). Erikson identified a range of psychosocial developmental stages from birth through to death. School-age children experience Erikson’s *4^th^ psychosocial stage*, described as a psychosocial crisis of industry vs inferiority. A child in this stage is often expected to learn and demonstrate new skills, productively complete tasks, and meet the expectations of their parents and teachers. During this stage, a child becomes aware of his/her abilities and the abilities of his/her peers. A child who cannot master these expected skills risks a sense of inferiority and failure. The potential impact of smartglasses on this developmental stage is not known. They may aid a child in successfully mastering this psychosocial stage by allowing him/her to be productive, and giving him/her a sense of achievement. There is also a risk that a child may feel inferior if s/he feels that without the smartglasses s/he is incompetent, or if s/he feels ridiculed for wearing such devices. Each individual child may face a unique situation based on his/her own personal attributes, and the support received from key people such as teachers, parents, and peers. This highlights the importance of ensuring that these key people are familiar with smartglasses technology, and understand its capabilities and functionality.

Following this stage is Erikson’s *5^th^ psychosocial stage* that occurs in adolescence, described as a psychosocial crisis between identity vs role confusion (55). Adolescence is a time of tremendous biological and psychological change (56), and during this stage individuals seek to define their role in the world, seeking to address the existential question, *who am I and what can I be?* Individuals will try to find likeminded social groups, focus on relationships with peers, and pursue sense of belonging. Many questions remain unanswered about how smartglasses may impact people with ASD during this stage, especially given the many social challenges people with ASD encounter during this transition from childhood to adulthood (57). How will ASD and these technologies define the individual? Will these technologies help individuals to find their purpose or hinder them? The impact of such technology may depend on smartglasses’ physical attributes, their impact on social relationships, or individual person characteristics (as discussed within the scope of the *bioecological model (53)*).

Learning happens continuously in childhood, and the use of smartglasses technology may provide a digital means of enabling learning to occur in *Vygotsky*’s Zone of Proximal Development (ZPD) (58). Vygotsky originally described his ZPD as being *“the distance between the actual development level as determined by independent problem solving and the level of potential development as determined through problem solving under adult guidance or in collaboration with more capable peers”* (59). These smartglasses designed as assistive technologies may allow children to undertake and learn tasks that they would have found impossible, or very difficult, to do independently. A child with ASD normally has a number of challenges in being in the ZPD, such as becoming overwhelmed with new experiences, struggling with transitions in environment or activities, and coping with sensory stimuli (19). Sensor-rich smartglasses may be of particular utility here in that they are able to monitor the behavioral and physiologic functioning of a child, detecting when they are under- or over- stimulated, and adapting the learning experience in real-time to keep a child engaged, and in the ZPD (13).

### Victimization, socialization, and the school environment

School-age children with ASD are at risk of being stigmatized (60) and being victims of bullying (61) for multiple reasons. They have different developmental trajectories that may put them at greater risk of victimization than their neurotypically developing peers, especially when they have challenges in social skills and communication (61). They may struggle to recognize social cues and develop relationships with their peers, impeding their ability to be better integrated by the community (62–64). Bullying may be particularly problematic at school, where academic and social factors may be a source of considerable stress, anxiety, and mental health concerns in children (65–67). A school represents not only an academic establishment, but a complicated and highly social environment. Children in schools often balance interpersonal relationships with peers and staff, complex social hierarchies, and school rules that can dictate the most basic elements of children’s day (whom to play with, where to sit, and when to talk to others (62–64, 68)). Some reports have suggested that children with ASD have inherently low motivation or desire to join social groups, but recent evidence indicates this is not the case and many have a strong desire for acceptance (69–71). Therefore, it is important to consider the acceptability and design of any assistive device in the population, given the risk of stigma and social isolation (26). This is especially true for a device that is worn on the face.

### Glass – a new generation of assistive reality smartglasses

On July 18, 2017, X (a subsidiary of Alphabet Inc., formerly known as Google X) released a successor to Google Glass, one of the most recognizable wearable devices in the world (72). Glass is a head-mounted, wearable computer that has demonstrated utility in a variety of situations where operating a computer hands-free and while heads-up is of particular advantage^i^. Glass has also been developed as a technology that can deliver social and cognitive skills coaching to children and adults with ASD (13). To our knowledge, we have published the first studies of ASD-related software on the original edition of Glass, Glass Explorer Edition (12, 13, 16, 17) and here we present the first scientific study of the newly announced successor version, known as Glass Enterprise Edition.

In this context, the announcement of a major new development in head-worn computing from X (72) signaled a potentially major advance for assistive technology targeting populations who traditionally face significant social challenges (18). It also signaled that head-worn computer devices will continue to exist and to evolve. Many have wondered if Glass and the smartglasses device category would die away, in part because of perceptions around desirability and social acceptability of wearing this new category of device in public (32, 73). The several-year quiet period in news about Glass also caused some to worry that the entire line of assistive research would be stranded. Therefore, public backing and leadership from one of the largest companies in the world (72), in this case the inventor of the product (74), provides assurance that head-worn computer platforms will persist. Thus, the recent announcement of Glass Enterprise Edition suggests that the complex and time-consuming process of developing assistive software applications that use the unique features of these platforms is wise and likely to continue.

It would seem that Glass Enterprise Edition (which has updates to the form factor, usability, central processor, display, audio system, and other features) would represent a substantial advantage for assistive technology apps and algorithms for ASD. However, it remains unknown whether people with ASD would actually desire to wear the new device. We have previously shown in children and adults with ASD that assistive applications running on the original Glass device were tolerable (17), safe (15), and can temporarily reduce some symptoms associated with ASD (13, 16). However, small changes in devices can greatly affect the desire of potential users to wear them.

We gave eight children with ASD an opportunity to try Glass Enterprise Edition in a controlled, recorded environment, and to explore its features, usability, and visual characteristics. We observed and recorded the interaction of the children with the device. We then asked them if they would use it for one hour a day at their current school as an assistive device. We also conducted a post-session semi-structured interview with their caregivers, who accompanied the child and observed the whole session. We recruited children who had previously used a wearable social-emotional artificial-intelligence aid based on the original Glass Explorer Edition, so they were familiar with the concepts involved and could evaluate the new hardware. Our sample represented a broad age range and severity spectrum of ASD.

## METHODS

### IRB Statement

The use of the Brain Power Autism System running on multiple head-worn computing devices by children and adults with autism was approved by Asentral, Inc., Institutional Review Board, an affiliate of the Commonwealth of Massachusetts Department of Public Health.

### Participants

Eight children with clinically diagnosed ASD were asked to comment on and test the comfort, usability, and feasibility of Glass Enterprise Edition. The participants represented a wide range of school-aged children, ages 6.7 to 17.2 years (mean ± SD: 11.7 ± 3.3 years), included seven males and one female.

Participants were recruited via a web-based research interest form and had each participated in at least one session of an ongoing study of autism assistive apps, the Brain Power Autism System, running on the original version of Google™ Glass, officially known as Glass Explorer Edition. At the time of testing Glass Enterprise Edition, they had access to additional devices.

Caregivers rated the participants’ level of overall ASD functioning according to a subjective 7-point scale (1 = lowest-functioning / severe; 7 = highest-functioning / mild). Caregivers also rated verbal functioning on a similar scale (1 = fully non-verbal, 7 = fully conversational). Participants represented a large range of both overall ASD functioning (range 4 – 7 out of 7; mean ± SD = 5.6 ± 1.1) and verbal functioning (range 4 – 7 out of 7; mean ± SD = 5.5 ± 1.3).

### Data Collection Procedure

Participants were orientated to the testing procedure and had to opportunity to use the Glass Enterprise Edition device in a quiet, controlled room (the ‘testing room’). Following device exposure, participants and caregivers were interviewed about their experience in a separate room (the ‘interview room’). In the testing room the participant was seated and allowed to use the device while being observed by and interacting with his/her caregiver, who sat opposite. Study staff stood and/or sat nearby observing from the side as the participant interacted with the device and with the caregiver. Participants were able to handle and wear the smartglasses for several minutes, and to ask questions of the study staff. They were encouraged to explore the device and assess its comfort, look, convenience, style, and related factors. They were provided with any assistance they required to properly place the device on their heads and align it with their eyes, though little assistance was needed. Testing sessions were recorded via video and photographs. All participants and/or caregivers gave written consent for their images and video to be used in any manner.

Following the testing session, participants were led from the testing room to the interview room, and were asked questions in a semi-structured interview in the presence of their caregivers. Participants were asked to compare their experience with Glass Enterprise Edition to previous experiences with wearable assistive devices and gamified applications related to ASD.

Participants were then asked whether they would consider wearing and using the device for one hour each day. The Glass Enterprise Edition device was then shown to the child again, and specific features about the device were demonstrated, including the ability for the device to be folded (the original edition did not fold). The child was then given the device, and could play freely with it as they were asked questions including, “Would you wear this for one hour each day at school?” and “Was it fun to wear this?”. The caregiver was also interviewed, and asked questions including “Did the experience go better than you anticipated?”.

For participants with moderate verbal skills, questions were repeated several times, slowly. Participants were allowed to gesture and to vocalize freely, and were eventually asked until they verbalized a “yes” or “no”. Study staff interacted with the child in front of the caregiver, and continued until assured based on the caregiver interaction, that the questions were understood, considered, and accurately answered.

### Exclusions

Individuals who had a known history of epilepsy or seizure disorder were not asked to take part in this study. Users who had any uncontrolled or severe medical or mental health condition that would make participation in the study predictably hazardous were also not invited to participate.

## RESULTS

All eight children, who represented the full range of school ages (6 – 17 years old), successfully wore, interacted with, and explored one or more Glass Enterprise Edition devices (**Figure 2**). The devices were loaded with a suite of assisted-reality apps for social-emotional learning and self-coaching related to brain-based challenges and needs, as discussed elsewhere (13). Participants explored the devices for several minutes at their leisure, putting them on and taking them off, exploring the style, size, weight, and shape, features such as foldability; and speaking out loud in some cases (more verbal children) about their observations and questions. All children successfully transitioned to the interview room, where they responded to questions by the experimenter, accompanied and assisted by their caregivers as needed.

**Figure 2(A-H).**
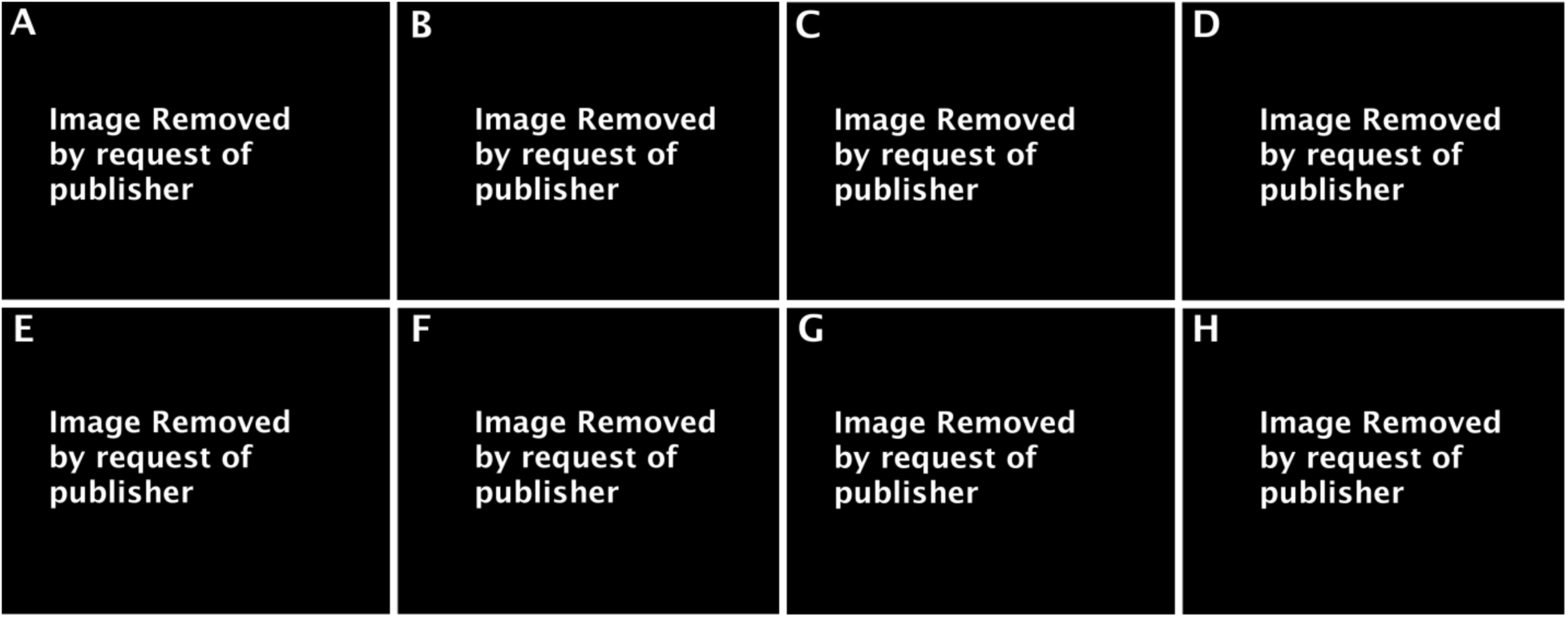
Children on the autism spectrum using and exploring the Glass Enterprise Edition device during a testing session at Brain Power. Each of the eight participants, who represented the entire range of school ages, and moderate to mild autism severity as well as moderately non-verbal to fully verbal functioning, rated Glass Enterprise Edition as desirable to wear on their heads and use daily in the often complex social environment of school. Glass Enterprise Edition was announced in the same month as the initial submission of this manuscript.

### Would you wear Glass Enterprise Edition for one hour each day at school?

Within the semi-structured interview following the testing session, each child with ASD was asked if s/he would wear Glass Enterprise Edition, as an assistive device, for one hour each day at her/his school. Each child was deemed by the study staff, based on verbal response and in conjunction with the caregiver’s comments and feedback, to have understood the question. Study staff was satisfied the children contemplated the question in the context of school and its social dynamics.

All eight children asserted that they would wear and use Glass Enterprise Edition at their schools (*n* = 8/8, 100%; **Table 1**).

**TABLE 1.**
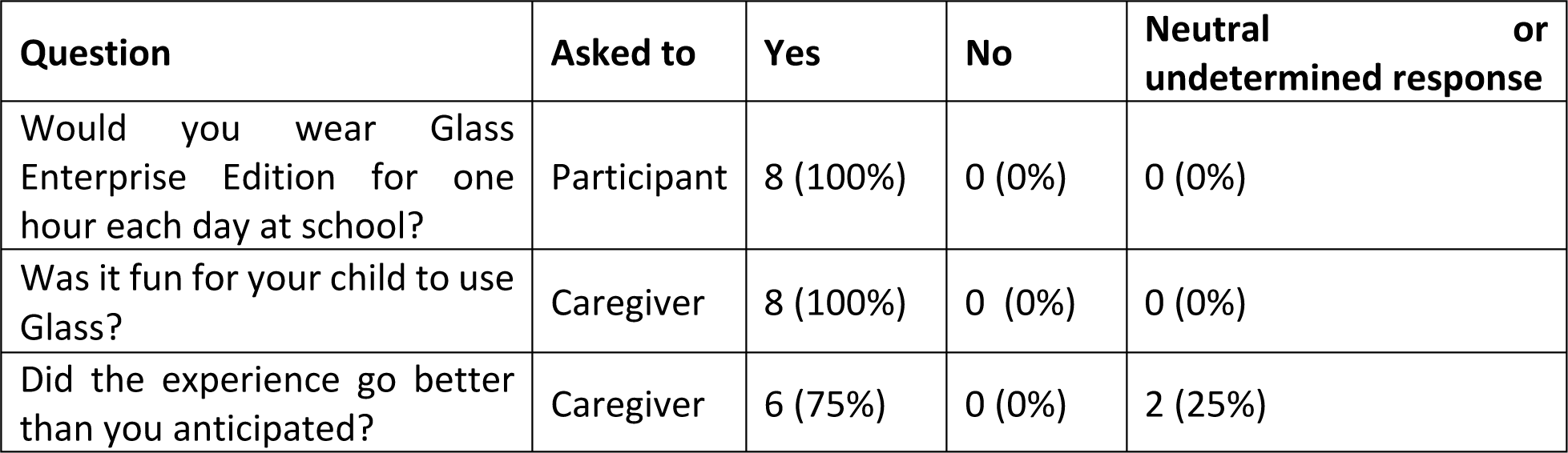

### Was it fun for your child to use Glass Enterprise Edition?

Each parent was asked to comment on whether it appeared to be fun for her/his child to test and explore the new device.

All eight parents responded that their children seemed to have fun during the experience (*n* = 8/8, 100%; **Table 1**).

### Did the experience go better than you anticipated?

Following each testing session, the caregiver was asked during a semi-structured interview whether s/he felt the experience went better than anticipated.

Of the eight parents, six parents asserted that the experience went better than they had thought, with one qualifying that it had gone “somewhat” better (*n* = 6/8, 75%; **Table 1**).

Of the remaining two, one parent said that the experience had proceeded “as expected”, and another answered the question conversationally but without a direct response, so the response was not tabulated as a “yes” but as an undetermined.

## Discussion

It is prudent to seek the opinions of children with ASD and their caregivers when considering the use of a new assistive device. Children with ASD and their caregivers may be particularly discerning about factors that could impact the use and social acceptance of such technologies in educational settings such as schools. Our study investigated the user acceptability of the updated version of Google Glass, known as Glass Enterprise Edition, a technology that was publicly announced by X the month of the initial submission of this manuscript. The eight school-aged children with ASD in this study unanimously rated Glass Enterprise Edition as an acceptable assistive technology for them to wear at school.

The current manuscript represents the first published work, to our knowledge, using Glass Enterprise Edition. It also represents the first published use of Glass Enterprise Edition as an assistive or assessment device for people with different abilities or intellectual disabilities or challenges. This work extends our previous research, which was the first peer-reviewed publication on the use of original Google Glass as an aid to people with ASD (13).

The results demonstrate that Glass Enterprise Edition was desirable by all participants, who spanned the full range of school ages (6 – 17 years old). However, the desirability in this case was predicated on a prediction of social acceptability (colloquially, the “cool factor”) in a social situation. Many factors may be included in a participant’s prediction of the cool factor of a device. Such factors may include unobtrusiveness, lightness, futuristic look, comfort, ease of storing, ease of transport, durability, ruggedness, styling, ability to give others experiences they could not otherwise have (conferring to the child an ability to control a social situation in a positive way), ability to initiate a conversation with decreased anxiety over selecting the topic of the conversation (“ice-breaker”), and more.

These results are important for a number of reasons. Children with ASD are frequently not involved in providing design or usability feedback to interventions and technologies that are developed for them. Involving children when choosing an assistive device is crucial to ensure that the device is socially appropriate for the environment, which will likely lead to greater compliance in wearing the device. It also appears that these children are accepting of new technologies, even on relatively uncommon and highly visible platforms, such as head-mounted computers. The children that participated in this study were more open to using Glass in a public environment than many adults have been (32). With this in mind, it will be equally as important to ensure caregivers and peers in the child’s *microsystem* are accepting of the assistive technology (54), as their opinions will likely sway a child’s enthusiasm towards the device. Many children in this study mentioned favoring Glass Enterprise Edition because of its unobtrusive, sleek design; having a device that is less noticeable and designed to be “cool” may help with its social acceptance and may not carry the stigma of assistive technology with it.

These results suggest that this platform may be suitable as a base for assistive software applications that could promote self-sufficiency. For instance, they may have a desirable new platform for gamified, social-emotional self-coaching applications based in neuroscience and artificial intelligence that have been deployed on other head-worn computer platforms (13). The results are promising at a broader level for those who wish to use or develop applications that harness the unique features of this family of devices, such as their ability to allow the user to be heads-up, hands-free, and able to perceive and engage with the world around while receiving additional assistance. Namely, the results suggest that the newest entrant into the still-emerging family of devices may be well received, at least by some discerning populations. Further research is clearly needed to address these and more limitations or open questions of the present work. This report represents a part of a larger, ongoing research initiative.

School is a place of high-risk relative to social integration, and stigma that could result from an undesirable or socially inappropriate device or behavior. This is one reason we chose the question of acceptability of the device at school as a high-bar test for how desirable and acceptable this new device may be. However, another limitation of the present work is that we asked for the opinion of the target users, and such an opinion is necessarily based on a prediction. It may be hard to predict how a device or behavior will actually be received in the complex and changing social hierarchy of a school environment. Additionally, children with ASD may have additional challenges in predicting the emotional reactions and behaviors of their classmates, especially if they are in an integrated school environment with neurotypical or typically-developing children their same chronological age. For all these reasons, further research is needed to test the acceptability within school environments.

